# Generation and Characterization of recombinant SARS-CoV-2 expressing reporter genes

**DOI:** 10.1101/2020.11.16.386003

**Authors:** Kevin Chiem, Desarey Morales Vasquez, Jun-Gyu Park, Roy Neal Platt, Tim Anderson, Mark R. Walter, James J. Kobie, Chengjin Ye, Luis Martinez-Sobrido

## Abstract

The emergence of severe acute respiratory syndrome coronavirus 2 (SARS-CoV-2), the pathogen responsible of coronavirus disease 2019 (COVID-19), has devastated public health services and economies worldwide. Despite global efforts to contain the COVID-19 pandemic, SARS-CoV-2 is now found in over 200 countries and has caused an upward death toll of over 1 million human lives as of November 2020. To date, only one Food and Drug Administration (FDA)-approved therapeutic drug (Remdesivir) and a monoclonal antibody, MAb (Bamlanivimab), but no vaccines, are available for the treatment of SARS-CoV-2. As with other viruses, studying SARS-CoV-2 requires the use of secondary approaches to detect the presence of the virus in infected cells. To overcome this limitation, we have generated replication-competent recombinant (r)SARS-CoV-2 expressing fluorescent (Venus or mCherry) or bioluminescent (Nluc) reporter genes. Vero E6 cells infected with reporter-expressing rSARS-CoV-2 can be easily detected via fluorescence or luciferase expression and display a good correlation between reporter gene expression and viral replication. Moreover, rSARS-CoV-2 expressing reporter genes have comparable plaque sizes and growth kinetics to those of wild-type virus, rSARS-CoV-2/WT. We used these reporter-expressing rSARS-CoV-2 to demonstrate their feasibility to identify neutralizing antibodies (NAbs) or antiviral drugs. Our results demonstrate that reporter-expressing rSARS-CoV-2 represent an excellent option to identify therapeutics for the treatment of SARS-CoV-2, where reporter gene expression can be used as valid surrogates to track viral infection. Moreover, the ability to manipulate the viral genome opens the feasibility of generating viruses expressing foreign genes for their use as vaccines for the treatment of SARS-CoV-2 infection.

**Importance:** Severe acute respiratory syndrome coronavirus 2 (SARS-CoV-2), the pathogen that causes coronavirus disease 2019 (COVID-19), has significantly impacted the human health and economic status worldwide. There is an urgent need to identify effective prophylactics and therapeutics for the treatment of SARS-CoV-2 infection and associated COVID-19 disease. The use of fluorescent- or luciferase-expressing reporter expressing viruses has significantly advanced viral research. Here, we generated recombinant (r)SARS-CoV-2 expressing fluorescent (Venus and mCherry) or luciferase (Nluc) reporter genes and demonstrate that they represent an excellent option to track viral infections *in vitro.* Importantly, reporter-expressing rSARS-CoV-2 display similar growth kinetics and plaque phenotype that their wild-type counterpart (rSARS-CoV-2/WT), demonstrating their feasibility to identify drugs and/or neutralizing antibodies (NAbs) for the therapeutic treatment of SARS-CoV-2. Henceforth, these reporter-expressing rSARS-CoV-2 can be used to interrogate large libraries of compounds and/or monoclonal antibodies (MAb), in high-throughput screening settings, to identify those with therapeutic potential against SARS-CoV-2.

## Introduction

Late in 2019, a previously unknown coronavirus, severe acute respiratory syndrome coronavirus 2 (SARS-CoV-2), was identified in Wuhan, China (1). Since then, SARS-CoV-2 has become responsible for the global pandemic of coronavirus disease 2019 (COVID-19) (1). As of November 2020, SARS-CoV-2 has spread worldwide and it has been responsible of over 40 million confirmed cases and around 1.1 million deaths (2). To date, only one United States (US) Food and Drug Administration (FDA)-approved therapeutic antiviral drug, Remdesivir, and a monoclonal antibody, MAb (Bamlanivimab) are available for the treatment of SARS-CoV-2 infections (3). No FDA-approved prophylactics (vaccines) are currently available against SARS-CoV-2.

SARS-CoV-2 is a single-stranded, positive-sense RNA *Betacoronavirus* that belongs to the *Coronaviridae* family. Prior to SARS-CoV-2, only six coronavirus (CoVs) species were known to cause disease in humans (4). Of the six, four human (h)CoVs are prevalent and responsible of causing common cold in immunocompetent individuals (hCoV-229E, hCoV-OC43, hCoV-NL63, and hCoV-HKU1) (4, 5). The two other CoVs, severe acute respiratory syndrome coronavirus (SARS-CoV) and Middle East respiratory syndrome coronavirus (MERS-CoV), have been associated with severe illness and significant morbidity and mortality (6). SARS-CoV was responsible for an outbreak of severe acute respiratory syndrome in 2002-2003 in Guangdong Province, China, with a fatality rate of around 9.5% (7). MERS-CoV was responsible for an outbreak of severe respiratory disease in 2012-2013 in the Middle East, with a fatality rate of around 30% (4, 5, 8). SARS-CoV-2 has a viral genome of approximately 30,000 nucleotides in length and high similarity to that of SARS-CoV (~79%) and lower to MERS-CoV (~50%), with an overall fatality rate of 3.4%, but can as high as 49% in critically ill patients, making the COVID-19 pandemic rival that of the “Spanish flu” in 1918-1919 (9–13).

Studying SARS-CoV-2 in laboratories require the use of secondary approaches to identify the presence of virus in infected cells. The ability to generate recombinant viruses using reverse genetics approaches represents a powerful tool to answer important questions in the biology of viral infections, including mechanisms of viral infection, pathogenesis and disease. In addition, the use of reverse genetics techniques have offered the possibility to generate recombinant viruses expressing reporter genes for their use in cultured cells or *in vivo* models of infection where reporter gene expression can be used as a valid surrogate the identify the presence of the virus in infected cells (14, 15). Importantly, these reporter-expressing recombinant viruses also represent an excellent tool for the easy and rapid identification of drugs for the prophylactic or therapeutic treatment of viral infections, by allowing high-throughput screening (HTS) approaches to interrogate large libraries of biologicals exhibiting antiviral activity.

Several manuscripts have described the ability to generate recombinant (r)SARS-CoV-2 expressing fluorescent (mNeonGreen and GFP) or bioluminescent (Nluc) reporter genes (16–18). However, these reverse genetics protocols require laborious *in vitro* assembly and transcription steps prior to transfecting cells, an inconvenience that should be considered due to the laborious nature and restraint of these methods. Here, we describe the generation and characterization of replication-competent rSARS-CoV-2 expressing fluorescent Venus or mCherry, or bioluminescent Nluc reporter genes using our recently described bacterial artificial chromosome (BAC)-based reverse genetics approach (19, 20). In Vero E6 cells, rSARS-CoV-2 expressing reporter genes have similar growth kinetics and plaque phenotype than that of wild-type virus (rSARS-CoV-2/WT). Importantly, we have observed a correlation between reporter gene expression and viral replication (19), and infected cells can be easily detected, without the need of secondary approaches, based on reporter gene expression. Using these reporter-expressing rSARS-CoV-2, we have developed fluorescent-based microneutralization assays that can be used to identify neutralizing antibodies (NAbs) and/or antivirals. The neutralization titers and inhibitory activities of NAbs or antivirals, respectively, obtained in our reporter-based microneutralization assays were similar to those observed in classical microneutralization assays using rSARS-CoV-2/WT (21). These results demonstrate that our reporter-expressing rSARS-CoV-2 represent an excellent tool for studying the biology of the virus and for the identification of therapeutics for the treatment of SARS-CoV-2 and also for *in vivo* studies. Furthermore, because of reporter gene expression, these rSARS-CoV-2 expressing reporter genes represent an ideal option to screen large libraries of biologicals to identify those with antiviral activity. Our results also demonstrate the feasibility of generating rSARS-CoV-2 expressing foreign genes that could be used to generate vaccines for the treatment of SARS-CoV-2 infections and/or associated COVID-19 disease.

## Materials and Methods

### Biosafety

All experiments involving infectious SARS-CoV-2 were performed in a biosafety level 3 (BSL3) laboratory at the Texas Biomedical Research Institute. Protocols containing SARS-CoV-2 were approved by the Texas Biomedical Research Institute’s Institutional Biosafety Committee (IBC).

### Cell lines

African green monkey kidney epithelial cells (Vero E6, CRL-1586) were grown and maintained in Dulbecco’s modified Eagle’s medium (DMEM) supplemented with 10% fetal bovine serum (FBS) and 1% PSG (100 units/ml penicillin, 100 μg/ml streptomycin, and 2 mM L-glutamine), at 37°C with 5% CO_2_.

### Generation of pBeloBAC11-SARS-CoV-2 encoding reporter genes

The pBeloBAC11 plasmid (NEB) containing the entire viral genome of SARS-CoV-2 has been previously described (19, 22). Briefly, the entire genome sequence of SARS-CoV-2 USA/WA1/2020 (GenBank accession no. MN985325) was chemically synthesized (Bio Basic) in five fragments and cloned into pUC57 plasmids containing unique restriction sites. Silent mutations were introduced to the spike (S) and matrix (M) genes to remove BstBI and Mlul restriction sites, respectively, that were used for the assembly of the entire SARS-CoV-2 genome into the pBeloBAC11 plasmid. These nucleotide changes were also used as genetic markers to distinguish the natural USA/WA1/2020 and the recombinant SARS-CoV-2 (19). The five fragments containing the entire SARS-CoV-2 genome were assembled into the pBeloBAC11 using standard molecular biology techniques. To remove the 7a gene and introduce the Venus, mCherry, or Nluc reporter genes, the region flanking the 7a viral gene and each individual reporter genes were amplified by extension and overlapping PCR using specific oligonucleotides in a shuttle plasmid. The modified 7a viral genes were inserted into the pBeloBAC11 plasmid containing the remaining SARS-CoV-2 viral genome using BamHI and RsrII restriction sites to generate pBeloBAC11-SARS-CoV-2-del7a/Venus, pBeloBAC11-SARS-CoV-2-del7a/mCherry, and pBeloBAC11-SARS-CoV-2-del7a/Nluc for the rescue of rSARS-CoV-2-Venus, rSARS-CoV-2-mCherry and rSARS-CoV-2-Nluc, respectively. Plasmids and pBeloBAC11 constructs were validated by Sanger sequencing (ACGT Inc).

### Rescue of rSARS-CoV-2 expressing reporter genes

The rSARS-CoV-2/WT and rSARS-CoV-2 expressing reporter genes were rescued as previously described (19, 20). Briefly, confluent monolayers of Vero E6 cells (1.2 × 10^6^ cells/well, 6-well plate format, triplicates) were transfected, using lipofectamine 2000 (LPF2000, Thermo Fisher) with 4 μg/well of pBeloBAC11-SARS-CoV-2/WT, pBeloBAC11-SARS-CoV-2-del7a/Venus, - del7a/mCherry, or - del7a/Nluc plasmids. An empty pBeloBAC11 plasmid was included as internal control. At 14 h, transfection media was replaced with post-infection media (DMEM with 2% FBS) and, 24 h later, cells were scaled up into T75 flasks. At 72 h, P0 virus-containing tissue culture supernatants were collected and stored at −80°C. Viral rescues were confirmed by infecting fresh Vero E6 cells (1.2 × 10^6^ cells/well, 6-wel plates, triplicates) and assessing fluorescence or Nluc expression. P0 viruses were passaged three times and viral stocks were generated and titrated for *in vitro* experiments. Viral titers (plaque forming units per milliliter; PFU/ml) were determined by plaque assay in Vero E6 cells (1.2 × 10^6^ cells/well, 6-well plate format).

### Sequencing

Viral RNAs from Vero E6 cells (1.2 × 10^6^ cells/well, 6-well plate format) infected at multiplicity of infection (MOI) of 0.01 were extracted using TRIzol reagent (Thermo Fisher Scientific), according to the manufacturer’s specifications. Libraries were generated with a KAPA RNA HyperPrep kit, 100 ng of RNA, and 7 mM of adapter. The Illumina HiSeq × was used for sequencing. Raw reads were filtered using Trimmomatic v0.39 (23). SARS-CoV-2 templates were made for each reporter gene by modifying SARS-CoV-2 USA/WA1/2020 (Genbank Accession: MN985325.1). Modifications included deleting orf7a, adding T21895C and T26843A mutations, and inserting the appropriate reporter gene (Venus, mCherry, or Nluc) at pos 27937. Reads were mapped to the modified SARS-CoV-2 templates with Bowtie v2.4.1 (24), and the total genomic coverage was quantified using MosDepth v0.2.6 (25). Allele frequencies were estimated with LoFreq* v2.1.3.1 (26) and low frequency variants with less than a 100x read depth or a 1% minor allele frequency were eliminated. All sequence data has been deposited in the NCBI Short Read Archive (BioProject: PRJNA678001).

### RT-PCR

Total RNA from Vero E6 cells (1.2 × 10^6^ cells/well, 6-well plate format) mock- or virus-infected (MOI of 0.01) were extracted using TRIzol reagent (Thermo Fisher Scientific). Superscript® II Reverse Transcriptase (Invitrogen) and Expand high-fidelity PCR (Sigma-Aldrich) were used to synthesize and amplify the cDNAs, respectively, using primers specific for the viral nucleoprotein (NP) or ORF7a region; and Venus, mCherry, or Nluc.

### Immunofluorescence assays (IFA)

Confluent monolayers of Vero E6 cells (1.2 × 10^6^ cells/well, 6-well format, triplicates) were mock-infected or infected (MOI of 0.01) with rSARS-CoV-2 expressing Venus or mCherry, or rSARS-CoV-2/WT. At 48 h post-infection, cells were fixed with 10% neutral buffered formalin at 4°C for 16 h for fixation and viral inactivation, and permeabilized with phosphate-buffered saline (PBS) containing 0.5% (vol/vol) Triton X-100 for 5 min at room temperature. Cells were washed with PBS and blocked with 2.5% bovine albumin serum (BSA) in PBS for 1 h before incubation with 1 μg/ml of SARS-CoV anti-NP MAb 1C7 in 1% BSA in PBS for 1 h at 37°C. Cells infected with rSARS-CoV-2-Venus or -mCherry were washed with PBS and stained with either Alexa Fluor 594 goat anti-mouse IgG (Invitrogen; 1:1000) or fluorescein isothiocynate (FITC)-conjugated goat anti-mouse IgG (Dako; 1:200), respectively. Cell nuclei were stained with 4’’,6’-diamidino-2-phenylindole (DAPI, Research Organics). Representative images were captured using a fluorescence microscope (EVOS M5000 imaging system) at 20X magnification.

### Protein gel electrophoresis and Western blots

Vero E6 cells (1.2 x10^6^ cells/well, 6-well plate format, triplicates) were mock-infected or infected (MOI of 0.01) with rSARS-CoV-2/WT or rSARS-CoV-2 expressing Venus, mCherry, or Nluc. At 48 h post-infection, cells were lysed with 1X passive lysis buffer (Promega) and proteins were separated by denaturing electrophoresis in 12% SDS-polyacrylamide gels and transferred to a nitrocellulose membrane (Bio-Rad) with a Mini-Protean Tetra Vertical Electrophoresis Cell at 100V for 1 h at 4°C. Membranes were blocked in PBS containing 10% dried skim milk and 0.1% Tween 20 for 1 h and then incubated overnight at 4°C with the following primary MAbs or polyclonal antibodies (PAbs): SARS-CoV NP (mouse MAb 1C7; Dr. Thomas Moran, Icahn School of Medicine at Mount Sinai), Venus (rabbit PAb sc-8334; Santa Cruz Biotech.), mCherry (rabbit PAb; Raybiotech), and Nluc (rabbit PAb; Promega). A MAb against actin (MAb AC-15; Sigma) was included as a loading control. Primary antibodies bound to the membrane were detected using horseradish peroxidase (HRP)-conjugated secondary antibodies against mouse or rabbit (GE Healthcare). Proteins were detected by chemiluminescence using SuperSignal West Femto Maximum Sensitivity Substrate (Thermo Scientific) based on the manufacturer’s specifications and imaged in a ChemiDoc imaging system (Bio-Rad).

### Plaque assays and immunostaining

Confluent monolayers of Vero E6 cells (1.2 × 10^6^ cells/well, 6-well plate format, triplicates) were infected with WT or reporter-expressing rSARS-CoV-2 for 1 h at 37°C. After viral absorption, infected cells were overlaid with agar and incubated at 37°C for 72 h. Afterwards, cells were submerged in 10% neutral buffered formalin at 4°C for 16 h for fixation and viral inactivation, and then the agar overlays were gently removed. To observe Venus and mCherry fluorescence expression, PBS was added to each well and plates were imaged under a fluorescence microscope (EVOS M5000 imaging system). For immunostaining, plates were permeabilized with 0.5% Triton X-100 PBS for 10 min at room temperature, blocked with 2.5% BSA PBS for 1 h at room temperature, and then incubated at 37°C for 1 h using the anti-SARS 2 NP MAb 1C7. Plaques were developed for visualization using the Vectastain ABC kit and DAB HRP Substrate kit (Vector laboratories), in accordance to the manufacturer’s recommendations.

### Viral growth kinetics and titrations

Vero E6 cells (1.2 × 10^6^ cells/well, 6-well plate format, triplicates) were infected (MOI of 0.01) with rSARS-CoV-2/WT or rSARS-CoV-2 expressing Venus, mCherry, or Nluc. After viral adsorption for 1 h at 37°C, cells were washed with PBS, provided with fresh post-infection media, and then placed in a 37°C incubator with 5% CO_2_ atmosphere. At the indicated times post-infection (12, 24, 48, 72, and 96 h), cells were imaged for Venus or mCherry expression under a fluorescence microscope (EVOS M5000 imaging system). Viral titers in the tissue culture supernatants at each time point were determined by titration and immunostaining, as previously described, using the anti-SARS-CoV NP MAb 1C7. Nluc expression in tissue culture supernatants was quantified using Nano-Glo luciferase substrate (Promega) following the manufacturer’s recommendations. Mean values and standard deviation (SD) were determined using GraphPad Prism software (version 8.2).

### Reporter-based microneutralization assay for the identification of antivirals

Vero E6 cells (96-well plate format, 4 × 10^4^ cells/well, quadruplicates) were infected with ~100-200 PFU of rSARS-CoV-2/WT or rSARS-CoV-2 expressing Venus, mCherry, or Nluc for 1 h at 37°C. After viral adsorption, cells were washed and incubated in 100 μL of infection media (DMEM with 2% FBS) containing 3-fold serial dilutions (starting concentration of 50 μM) of Remdesivir, or 0.1% DMSO vehicle control, and 1% avicel (Sigma-Aldrich). Cells infected with rSARS-CoV-2/WT or rSARS-CoV-2 expressing fluorescent Venus or mCherry were incubated at 37°C for 24 h, while cells infected with rSARS-CoV-2 expressing Nluc were incubated at 37°C for 48 h. For rSARS-CoV-2/WT and rSARS-CoV-2 expressing fluorescent Venus and mCherry, cells were submerged in 10% neutral buffered formalin at 4°C for 16 h for fixation and viral inactivation. Cells were washed with 100 μl/well of PBS three times, permeabilized with 100 μl/well of 0.5% Triton X-100 in PBS at room temperature for 15 min and blocked with 100 μl/well of 2.5% BSA in PBS at 37°C for 1 h. Next, cells were staining with the anti-NP MAb 1C7 (1μg/mL) in 1% BSA PBS at 37°C for 1 h. After incubation with the primary MAb, cells were washed with PBS three times, and a secondary fluorescein isothiocynate (FITC)-conjugated goat anti-mouse IgG (Dako; 1:200) in 1% BSA were added to cells for 1 h at 37°C. Cell nuclei were stained with 4’’,6’-diamidino-2-phenylindole (DAPI, Research Organics). Viral infections were determined using fluorescent images of each well and quantified using a cell image analysis software, Cell Profiler (Broad Institute). In the case of cells infected with rSARS-CoV-2 expressing Nluc, tissue culture supernatants were collected at 48 h post-infection and Nluc expression was measured using a luciferase assay and a Synergy LX microplate reader (BioTek). Individual wells from three independent experiments conducted in quadruplicates were used to calculate the average and standard deviation (SD) of viral inhibition using Microsoft Excel software. Non-linear regression curves and the half maximal effective concentration (EC_50_) of Remdesivir was determined using GraphPad Prism software (version 8.2).

### Reporter-based microneutralization assay for the identification of NAbs

To test the neutralizing activity of 1212C2, a human MAb recently described to neutralize SARS-CoV-2 (27), confluent monolayers of Vero E6 cells (96-plate format, 4 × 10^4^ cells/well, quadruplicates) were infected with ~100-200 PFU of rSARS-CoV-2/WT or rSARS-CoV-2 expressing Venus, mCherry, or Nluc for 1 h at 37°C. After viral adsorption, cells were washed and incubated with 100 μL of infection media (DMEM 2% FBS) containing 3-fold serial dilutions (starting concentration of 500 ng) of 1212C2 or PBS, and 1% avicel (Sigma-Aldrich). Infected cells were incubated at 37°C for 24 h for rSARS-CoV-2/WT, or rSARS-CoV expressing Venus or mCherry, and 48 h for rSARS-CoV-2 expressing Nluc. After viral infections, cells infected with rSARS-CoV-2/WT and rSARS-CoV-2 expressing fluorescent Venus and mCherry were submerged in 10% neutral buffered formalin at 4°C for 16 h for fixation and viral inactivation. Cells were washed with 100 μl/well of PBS three times, permeabilized with 100 μl/well of 0.5% Triton X-100 in PBS at room temperature for 15 min. Then, cells were blocked with 100 μl/well of 2.5% BSA in PBS at 37°C for 1 h. Cells were next incubated with the anti-NP MAb 1C7 (1μg/ml) in 1% BSA PBS at 37°C for 1 h. Cells were next washed three times with PBS and incubated with a secondary fluorescein isothiocynate (FITC)-conjugated goat anti-mouse IgG (Dako; 1:200) in 1% BSA for 1 h at 37°C. Cell nuclei were stained with 4’’,6’-diamidino-2-phenylindole (DAPI, Research Organics). Viral infections were determined using fluorescent images of each well and quantified using a cell image analysis software, Cell Profiler (Broad Institute). In the case of cells infected with rSARS-CoV-2 expressing Nluc, tissue culture supernatants were collected at 48 h post-infection and Nluc expression was measured using a luciferase assay and a Synergy LX microplate reader (BioTek). Individual wells from three independent experiments conducted in quadruplicates were used to calculate the average and standard deviation (SD) of viral inhibition using Microsoft Excel software. Non-linear regression curves and the half maximal neutralizing concentration (NT_50_) of 1212C2 was determined using GraphPad Prism software (version 8.2).

### Genetic stability

Vero E6 cells (1.2 × 10^6^ cells/well, 6-well plate format, triplicates) were infected (MOI of 0.01) with rSARS-CoV-2-Venus or -mCherry P3 stocks and after 1 h viral adsorption, virus inoculum was replaced with infectious media (DMEM 2% FBS). The cells were incubated at 37°C with 5% CO_2_ until 70% cytopathic effect (CPE) was observed. Then, tissue culture supernatants were collected and diluted 100-fold in infectious media and used to infect fresh Vero E6 cells (1.2 × 10^6^ cells/well, 6-well format, triplicates) for two additional passages (P5). Venus- and mCherry-expressing plaques (~50 counted plaques per viral passage) were evaluated by immunostaining and fluorescent protein expression. Viral plaques were imaged under a fluorescence microscope (EVOS M5000 imaging system) under 4X magnification.

## Results

### Generation of rSARS-CoV-2 expressing reporter genes

The pBeloBAC11 plasmid encoding the full-length viral genome of SARS-CoV-2 was previously described (19). To generate the reporter-expressing rSARS-CoV-2, the 7a open reading frame (ORF) was substituted with Venus, mCherry, or Nluc gene in the pBeloBAC11 plasmid encoding the remaining viral genome to produce pBeloBAC11-SARS-CoV-2-del7a/Venus, -del7a/mCherry, or -del7a/Nluc plasmids for viral rescues. We then used our previously described BAC-based reverse genetics approach to rescue rSARS-CoV-2-Venus, -mCherry, and -Nluc (**Figure 1A**).

**Figure 1.**
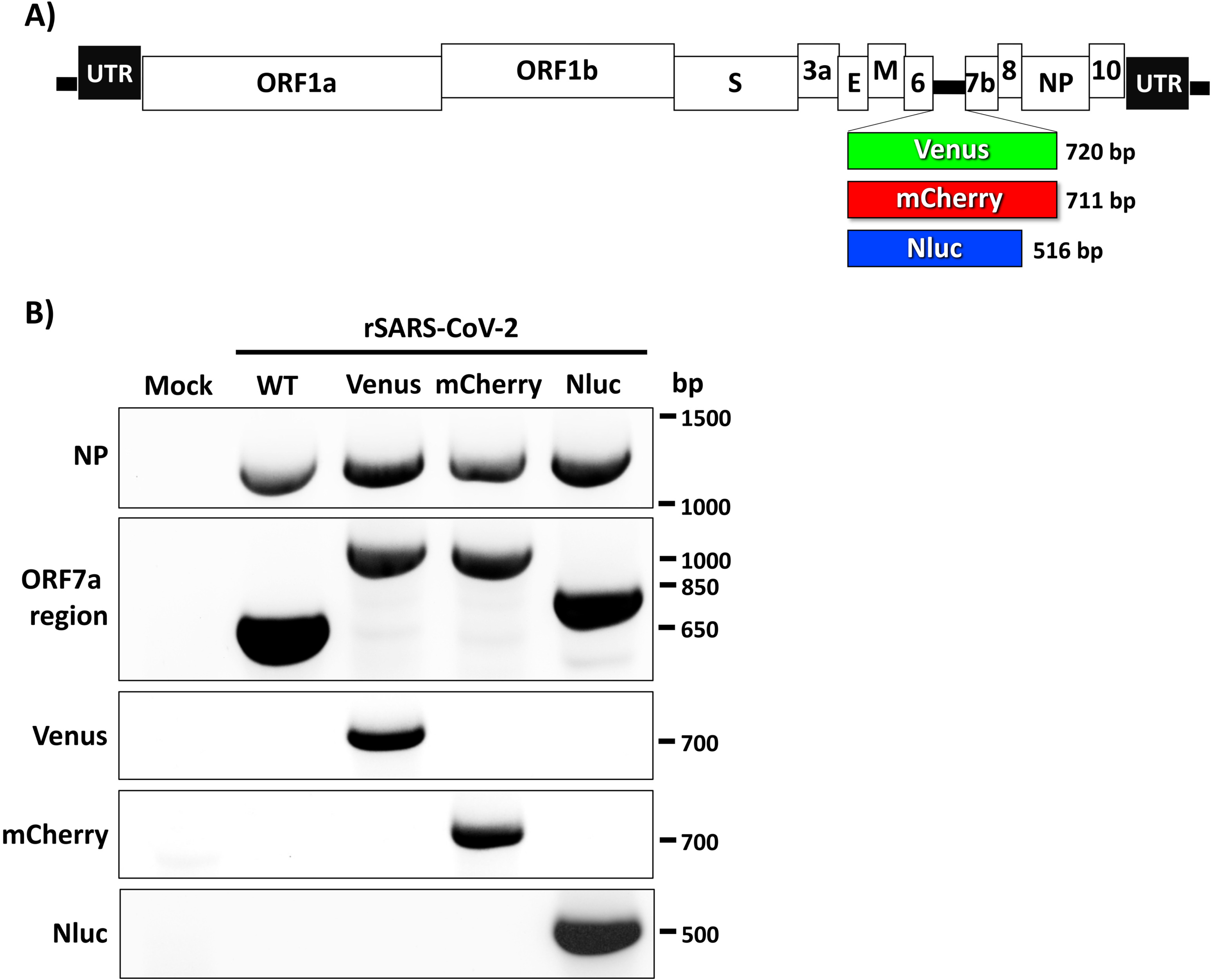
rSARS-CoV-2 expressing reporter genes. **A) Schematic representation of reporter-expressing rSARS-CoV-2:** rSARS-CoV-2 expressing Venus (green box), mCherry (red box), and Nluc (blue box) reporter genes instead of the viral 7a ORF are shown. The molecular size of the three reporter genes are indicated. The location of other viral proteins and untranslated regions (UTR) are also shown. **B) Genetic characterization of reporter-expressing rSARS-CoV-2:** Vero E6 cells were mock-infected or infected (MOI 0.01) with WT or reporter-expressing rSARS-CoV-2. At 72 h post-infection, total RNA collected from cells was used to amplify, using RT-PCR, the viral NP, the ORF 7a region, and the different reporter genes (Venus, mCherry or Nluc). Primers used for this RT-PCR analysis are shown in the left. The molecular weight (bp) of the RT-PCR amplified products is shown on the right.

We confirmed the rescue of rSARS-CoV-2 expressing -Venus, -mCherry, or -Nluc reporter genes by RT-PCR using total RNA from mock-, rSARS-CoV-2/WT- or rSARS-CoV-2 reporter virus-infected cells using primers specific for the viral NP, the ORF7a region, or the individual reporter genes (**Figure 1B**). As expected, primers specific for SARS-CoV-2 NP amplified a band of ~1260 bp from the RNA extracted from rSARS-CoV-2-infected but not mock-infected cells (**Figure 1B**). Amplified bands using primers in the ORF7a region resulted in the expected ~566 bp in cells infected with rSARS-CoV-2/WT and ~ 920, 911, and 815 bp in the case of cells infected with rSARS-CoV-2-Venus, -mCherry and -Nluc, respectively, based on the different size of the reporter genes (**Figure 1B**). Primers specific for the reporter genes only results in the RT-PCR amplification of bands from cells infected with the respective reporter-expressing rSARS-CoV-2 (**Figure 1B**). These results demonstrate that substitution of the viral ORF7a for Venus, mCherry, or Nluc genes results in the successful recovery of rSARS-CoV-2 containing these reporter genes.

### Characterization of rSARS-CoV-2 expressing reporter genes

Next, we characterize the reporter-expressing rSARS-CoV-2 by evaluating the expression levels of Venus, mCherry, or Nluc in cell cultures, and compared them to those of cells infected with rSARS-CoV-2/WT (**Figure 2**). The rSARS-CoV-2 expressing Venus and mCherry were directly visualized under a fluorescence microscope (**Figure 2A**). Indirect immunofluorescence microscopy using a MAb against SARS-CoV NP was used to detect rSARS-CoV-2/WT infection (**Figure 2A**). As expected, Venus or mCherry expression were only observed in Vero E6 cells infected with rSARS-CoV-2 expressing Venus or mCherry, respectively, but not in cells infected with rSARS-CoV-2/WT (**Figure 2A**). Importantly, only cells infected with rSARS-CoV-2-Venus or rSARS-CoV-2-mCherry were detected using green or red filters, respectively (data not shown). As expected, the viral NP was detected in cells infected with rSARS-CoV-2-WT, -Venus, or -mCherry (**Figure 2A**). Expression of Nluc in rSARS-CoV-2-Nluc-infected cells was evaluated from tissue culture supernatants at 48 h post-infection (**Figure 2B**). High levels of Nluc expression were detected in culture supernatants of cells infected with rSARS-CoV-2-Nluc but not from mock or rSARS-CoV-2/WT infected cells (**Figure 2B**). These results demonstrate that Vero E6 cells infected with rSARS-CoV-2-Venus, -mCherry, or -Nluc expresses the corresponding reporter genes and that viral infections can be detected by fluorescence (rSARS-CoV-2-Venus or -mCherry) or luciferase (rSARS-CoV-2-Nluc) without the need of antibodies that were required for the detection of rSARS-CoV-2/WT.

**Figure 2.**
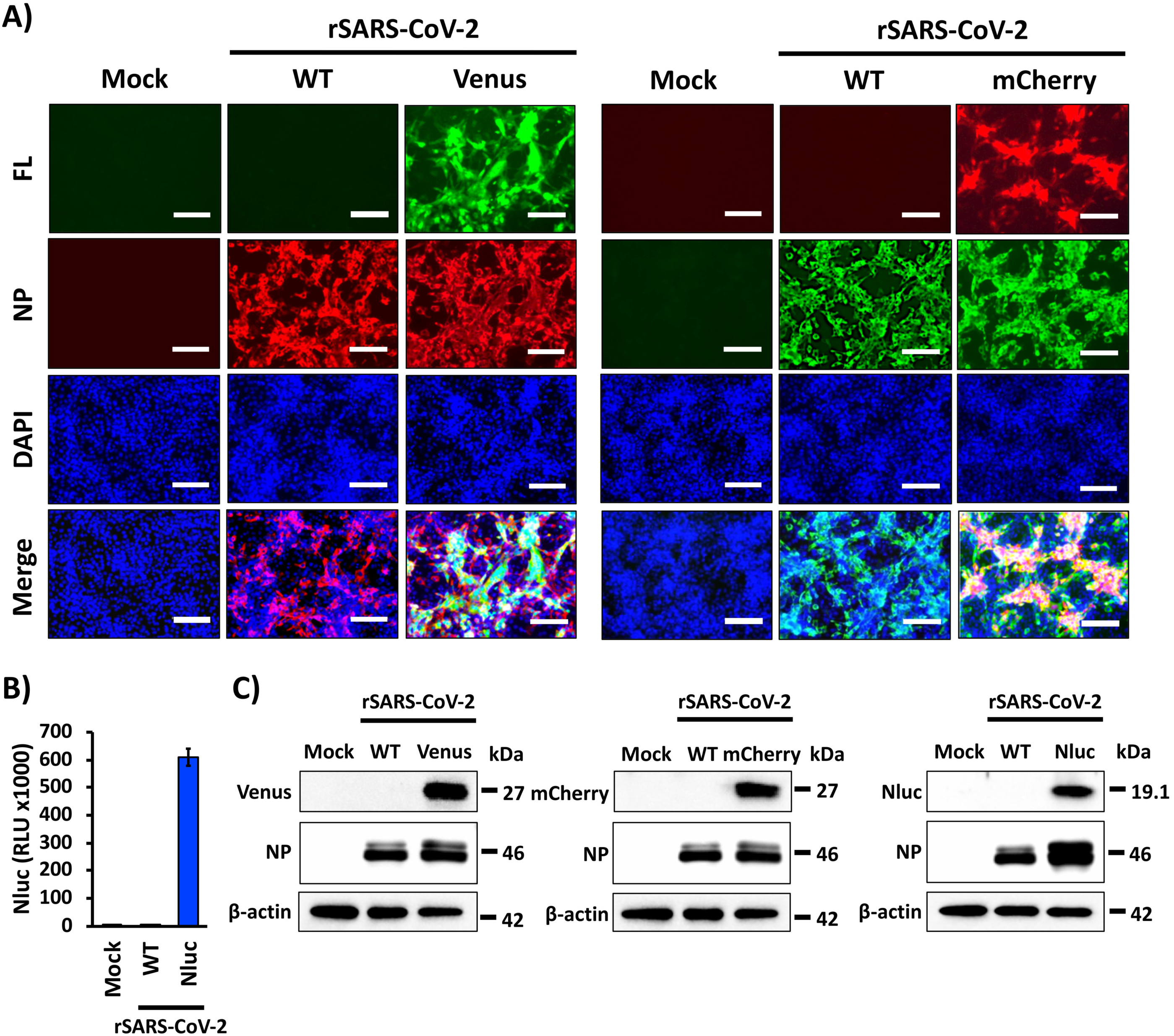
Characterization of reporter-expressing rSARS-CoV-2. **A) Fluorescent expression:** Vero E6 cells were mock infected or infected (MOI 0.01) with WT and Venus- or mCherry-expressing rSARS-CoV-2. At 48 h post-infection, cells were fixed and permeabilized, visualized for Venus (left) or mCherry (right) expression, and immunostained with a SARS-CoV NP MAb (1C7). DAPI was used for nuclear staining. Merged images for Venus (left) or mCherry (right), viral NP, and DAPI are illustrated. Representative images (20X magnification) are shown. Scale bar, 100 μm. **B) Nluc expression:** Vero E6 cells were mock-infected or infected (MOI 0.01) with WT and Nluc-expressing rSARS-CoV-2. At 48 h post-infection, Nluc expression in tissue culture supernatants was analyzed using a Synergy LX microplate reader (BioTek). **C) Western blot:** Vero E6 cells were mock-infected or infected (MOI 0.01) with WT and Venus (left), mCherry (center) or Nluc (right) expressing rSARS-CoV-2. At 48 h post-infection, viral NP and reporter gene protein expression levels were analyzed using specific antibodies. An antibody against beta-actin was used as internal control. The size of molecular markers is shown in the right in each of the Western blots.

We next evaluated reporter protein expression levels by Western blot assay using cell lysates from either mock, rSARS-CoV-2-WT, or rSARS-CoV-2-Venus, -mCherry, or -Nluc infected cells using MAbs against the viral NP, the reporter genes, or actin as a loading control (**Figure 2C**). As expected, reporter gene expression was detected in cell lysates of cells infected with the respective reporter-expressing rSARS-CoV-2 but not from mock or rSARS-CoV-2-WT infected cells. Viral NP expression was detected in cell lysates from all virus-infected cells, but not mock-infected cells (**Figure 2C**).

Next, we assessed reporter gene expression over a period of 96 h in cells that were mock-infected (data not shown) or cells infected with WT or reporter-expressing rSARS-CoV-2 (**Figure 3**). Venus and mCherry expression levels were determined using fluorescence microscope (**Figure 3A**), while Nluc activity in tissue culture supernatants from infected cells was detected using a luminometer (**Figure 3B**). Venus and mCherry expression were detected as early as 24 h post-infection and fluorescent protein expression increased over time until 96 h post-infection where a decrease in fluorescence was observed because of CPE caused by viral infection (brightfield, BF) (**Figure 3A**). Similar CPE, but not fluorescent expression, was also observed in cells infected with rSARS-CoV-2/WT (**Figure 3A**). Levels of Nluc expression were also detected as early as 24 h post-infection and increase in a time-dependent matter (**Figure 3B**).

**Figure 3.**
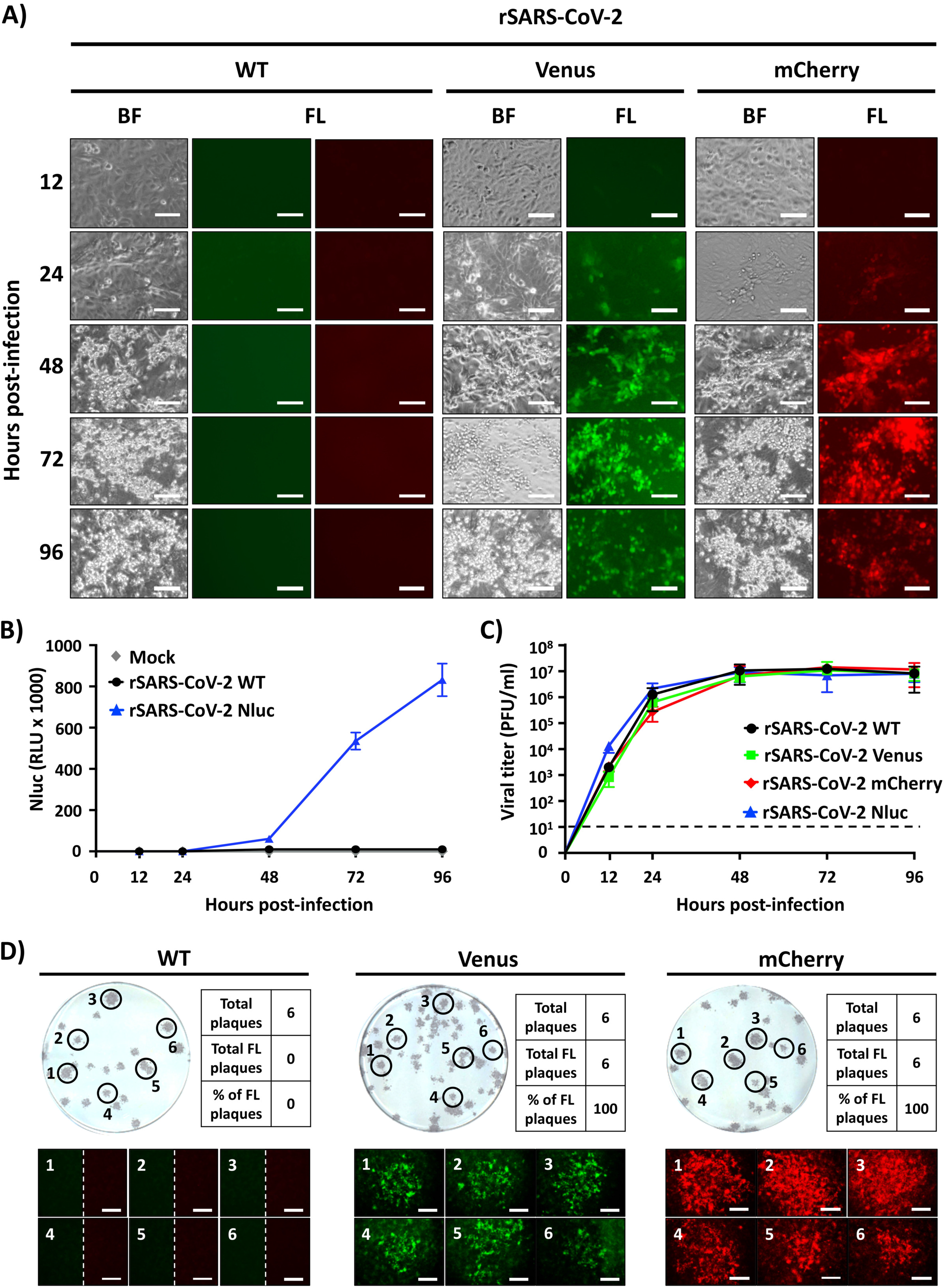
Viral growth kinetics and plaque phenotype. **A) Fluorescent expression:** Vero E6 cells were infected (MOI 0.01) with WT (left), Venus (center), and mCherry (right) expressing rSARS-CoV-2. At 12, 24, 48, 72, and 96 h post-infection. fluorescence protein expression was determined using a fluorescent microscope. Representative images (20X magnification) are included. Scale bar, 100 μm. **B) Nluc expression:**Vero E6 cells were mock-infected or infected (MOI 0.01) with WT and Nluc expressing rSARS-CoV-2. At the indicated times post-infection (12, 24, 48, 72, and 96 h), Nluc expression in the tissue culture supernatants was analyzed using a Synergy LX microplate reader (BioTek). **C) Growth kinetics:** Vero E6 cells were infected (MOI 0.01) with WT or reporter-expressing rSARS-CoV-2. At 12, 24, 48, 72, and 96 h post-infection, presence of infectious virus in the tissue culture supernatants was determined using plaque assay (plaque forming units, PFU/ml). **D) Plaque phenotype:** Vero E6 cells were infected with ~25 PFU of WT (left), Venus (middle), and mCherry (right) expressing rSARS-CoV-2. At 72 h post-infection, plaques were observed under a fluorescent microscope to detect Venus or mCherry expression. In the case of rSARS-CoV-2/WT infected cells, images correspond to fluorescent filters to detect Venus (left) or mCherry (right) expression. Thereafter, viral plaques were detected using the 1C7 SARS-CoV NP MAb. A selected number (n=6) of plaques were used to determine the percentage of viral plaques expressing fluorescent proteins (Venus or mCherry). Magnification 4x, Scale bar, 750μm.

To assess whether deletion of 7a ORF and insertion of reporter genes compromised viral fitness in cultured cells, we compared growth kinetics of reporter-expressing rSARS-CoV-2 to those of rSARS-CoV-2/WT (**Figure 3C**). We found all the reporter-expressing rSARS-CoV-2 exhibited similar growth kinetics and peak viral titers of infection to that of rSARS-CoV-2/WT (**Figure 3C**), suggesting that deletion of the 7a ORF and insertion of the reporter genes did not significantly affect viral fitness, at least in cultured cells. These results also support previous findings with SARS-CoV where deletion of the 7a ORF and insertion of reporter genes did not impact viral fitness *in vitro* (28, 29). These results were further confirmed when we evaluate the plaque phenotype of the rSARS-CoV-2 expressing fluorescent reporter genes and compared them to those of rSARS-CoV-2/WT (**Figure 3D**). Similar plaque sizes were observed in Vero E6 cells infected with rSARS-CoV-2/WT and rSARS-CoV-2 expressing Venus or mCherry (**Figure 3D**). Notably, Venus-positive or mCherry-positive plaques were only detected in cells infected with rSARS-CoV-2-Venus or -mCherry, respectively, and not in rSARS-CoV-2/WT infected cells (**Figure 3D**). Importantly, fluorescent plaques overlapped with those detected by immunostaining using the SARS-CoV NP 1C7 MAb. Similar to the growth kinetics data, we found no significant differences in the plaque size of reporter-expressing rSARS-CoV-2 compared to rSARS-CoV-2/WT (**Figure 3D**).

### A reporter-based microneutralization assay for the identification of antivirals

To determine the feasibility of using our reporter-expressing rSARS-CoV-2 for the identification of antivirals, we evaluated the ability of Remdesivir to inhibit SARS-CoV-2 in reporter-based microneutralization assays (**Figure 4**). Remdesivir has been previously described to inhibit SARS-CoV-2 infection and is the only FDA-approved antiviral for the treatment of SARS-CoV-2 (3, 21, 30). The EC_50_ of Remdesivir against rSARS-CoV-2-Venus (**Figure 4A**, 1.07 μM), -mCherry (**Figure 4B**, 1.78 μM), or Nluc (**Figure 4C**, 1.79 μM) were similar to those obtained with rSARS-CoV-2/WT (**Figure 4D**, 1.51 μM) and values previously reported in the literature (21). This demonstrates the feasibility of using these reporter-expressing rSARS-CoV-2 and the reporter-based assay to easily identify compounds with antiviral activity based on fluorescent or luciferase expression and without the need of MAbs to detect the presence of the virus in infected cells.

**Figure 4.**
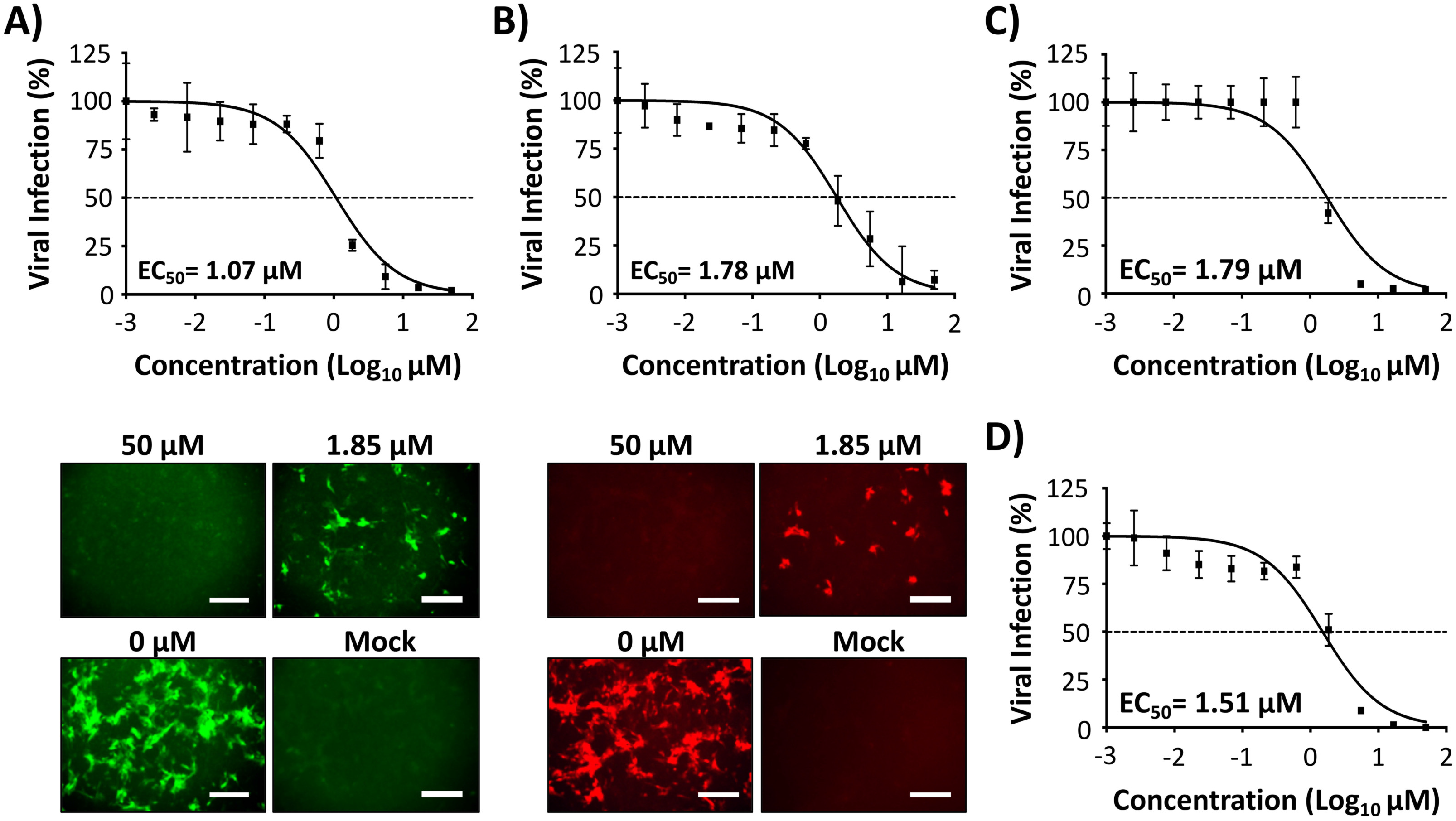
A reporter-based microneutralization assay for the identification of antivirals: Vero E6 cells (96-well plate format, ~4 × 10^4^ cells/well, triplicates) were infected with 100 PFU of Venus (**A**), mCherry (**B**), Nluc (**C**), or WT (**D**) rSARS-CoV-2. After 1 h viral absorption, post-infection media containing 3-fold serial dilutions of Remdesivir (starting concentration 50 μM) was added to the cells. At 24 h post-infection, cells were fixed and visualized for Venus (**A**) and mCherry (**B**) expression using a fluorescent microscope. In the case of cells infected with rSARS-CoV-2 expressing Nluc, luciferase expression in the tissue culture supernatant was determined at 48 h post-infection using a luciferase assay and a Synergy LX microplate reader (BioTek) (**C**). For the detection of rSARS-CoV-2/WT, the amount of virus was determined by plaque assay using the 1C7 SARS-CoV NP MAb (**D**). The amount of viral infection for Venus-, mCherry-, or WT-rSARS-CoV-2 (after IFA) was determined using fluorescent images of each well and quantified using a cell image analysis software, Cell Profiler (Broad Institute). Nluc activity was quantified using the Gen5 data analysis software (BioTek). The 50% effective concentration (CC_50_) of Remdesivir was determined using Graphpad Prism. Dotted line indicates 50% viral inhibition. Data were expressed as mean and SD from triplicate wells. Representative images (10X magnification) are included. Scale bar, 300μm.

### A reporter-based microneutralization assay for the identification of NAbs

We next evaluate the feasibility of using our reporter-expressing rSARS-CoV in a reporter-based microneutralization assay to identify NAbs against SARS-CoV-2. As proof of concept, we used a human MAb (1212C2) which we have recently described to potently bind and neutralize SARS-CoV-2 infection both *in vitro* and *in vivo* (27). The NT_50_ of 1212C2 against rSARS-CoV-2-Venus (**Figure 5A**, 1.94 ng), -mCherry (**Figure 5B**, 5.02 ng), or Nluc (**Figure 5C**, 3.67 ng) were similar to those observed with rSARS-CoV-2/WT (**Figure 5D**, 4.88 ng), and recently reported values (27).

**Figure 5.**
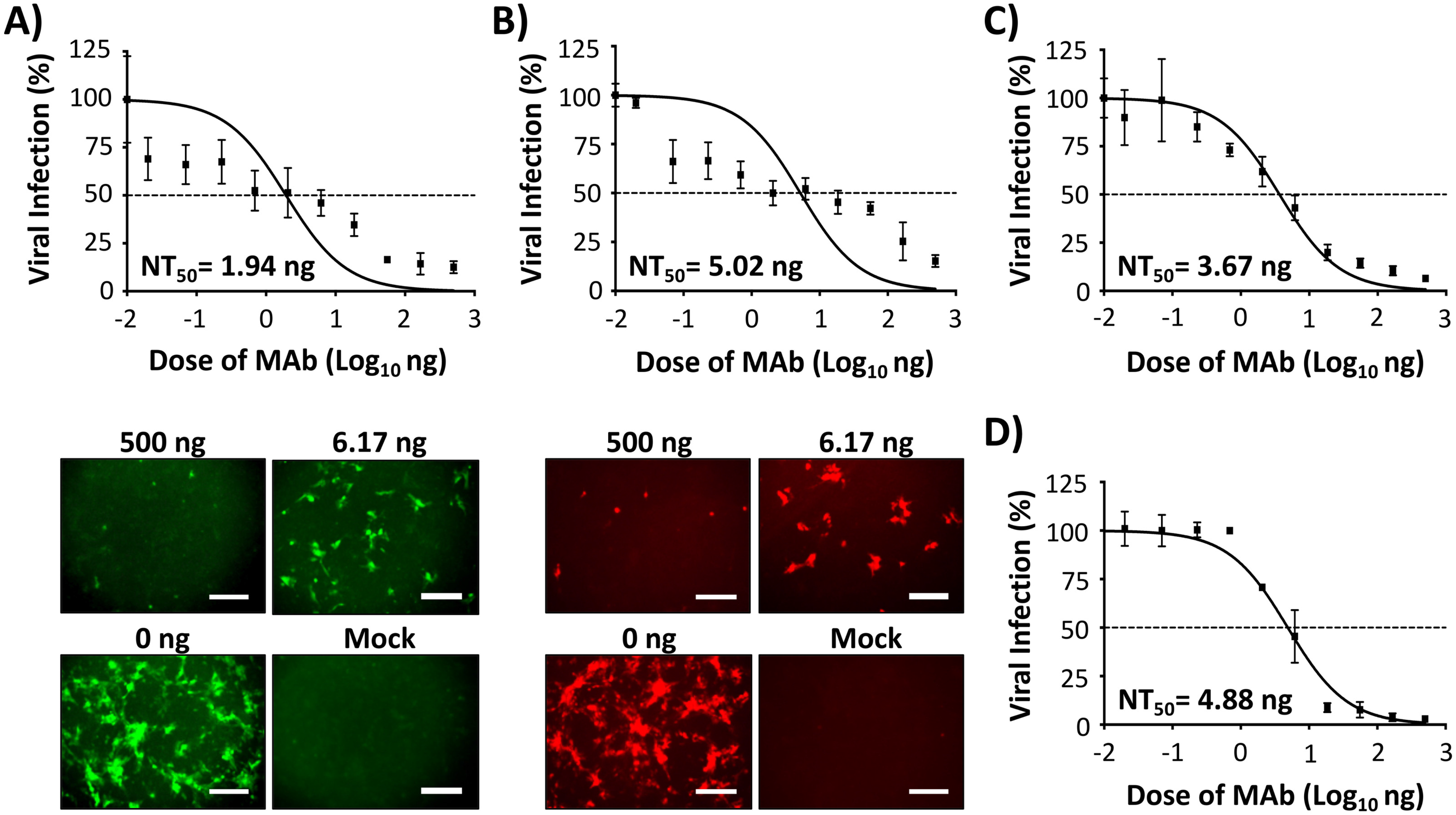
A reporter-based microneutralization assay for the identification of NAbs: Vero E6 cells (96-well plate format, ~4 × 10^4^ cells/well, triplicates) were infected with 100 PFU of Venus (**A**), mCherry (**B**), Nluc (**C**) or WT (**D**) rSARS-CoV-2. After 1 h viral absorption, post-infection media containing 3-fold serial dilutions (starting concentration 500 ng) of a SARS-CoV-2 NAb (1212C2) was added to the cells. At 24 h post-infection, cells were fixed and visualized for Venus (**A**) and mCherry (**B**) expression using a fluorescent microscope. In the case of cells infected with rSARS-CoV-2 expressing Nluc, luciferase expression in the tissue culture supernatant was determined at 48 h post-infection using a luciferase assay and a Synergy LX microplate reader (BioTek) (**C**). For the detection of rSARS-CoV-2/WT, the amount of virus was determined by plaque assay using the 1C7 SARS-CoV NP MAb (**D**). The amount of viral infection for Venus-, mCherry-, and WT-rSARS-CoV-2 (after IFA) was determined using fluorescent images of each well and quantified using a cell image analysis software, Cell Profiler (Broad Institute). Nluc was quantified using the BioTek Gen5 data analysis software. The 50% neutralizing titer (NT_50_) of 1212C2 was determined using Graphpad Prism. Dotted line indicates 50% viral neutralization. Data were expressed as mean and SD from triplicate wells. Representative images (10X magnification) are included. Scale bar, 300μm.

### Genetic stability of rSARS-CoV-2 *in vitro*

The genetic stability of reporter-expressing recombinant viruses is important to demonstrate their viability in *in vitro* and/or *in vivo* studies. To evaluate the ability of our rSARS-CoV-2 to maintain fluorescent reporter gene expression, viruses were consecutively passaged in Vero E6 cells and Venus and mCherry expression were determined by plaque assay using fluorescent microscopy (**Figure 6A**). To that end, we evaluated fluorescent expression of over 40 plaques before immunostaining with an anti-SARS-CoV NP MAb 1C7. We found the Venus and mCherry fluorescent expression from our rSARS-CoV-2 was genetically stable with nearly 100% of the plaques analyzed under a fluorescent microscope (**Figure 6A**). We also evaluated the complete genome sequences of the reporter-expressing rSARS-CoV-2 used in our studies (P3) with those of additional passages (P4 and P5) using next generation sequencing (**Figure 6B**). In the case of rSARS-CoV-2/Venus (**Figure 6B, top**), few variants were found at low frequencies after two additional passages (P5), indicating no significant changes and/or deletions in the viral genome. However, for rSARS-CoV-2/mCherry (**Figure 6B, middle**), variants containing a mutation at position 21,784 in the S gene was found in our viral stock (P3) and the frequency of this mutation increased after additional passages (P4 and P5). Two additional mutations at positions 23,525 and 24,134 (bot in the S gene) were also found at P5. In the case of rSARS-CoV-2/Nluc (**Figure 6B, bottom**) a mutation at position 24,755 (S gene) was found in our viral stock (P3). Frequency of this mutation increased up to 100% after 2 additional passage (P5). Other, less abundant, mutations at positions 13,419 (nsp12, RNA dependent RNA polymerase), 23,525 (S gene), and 26,256 (envelop, E, gene) were also found after the additional 2 passages (P5) (**Figure 6B, bottom**). It is possible that these mutations are most likely due to viral adaptation to Vero E6 cells but since different mutations were found in the three reporter-expressing rSARS-CoV-2, it is also possible that these mutations are related to the nature of the reporter gene. In any case, these results indicate that the reporter-expressing rSARS-CoV-2 are genetically stable in Vero E6 cells.

**Figure 6.**
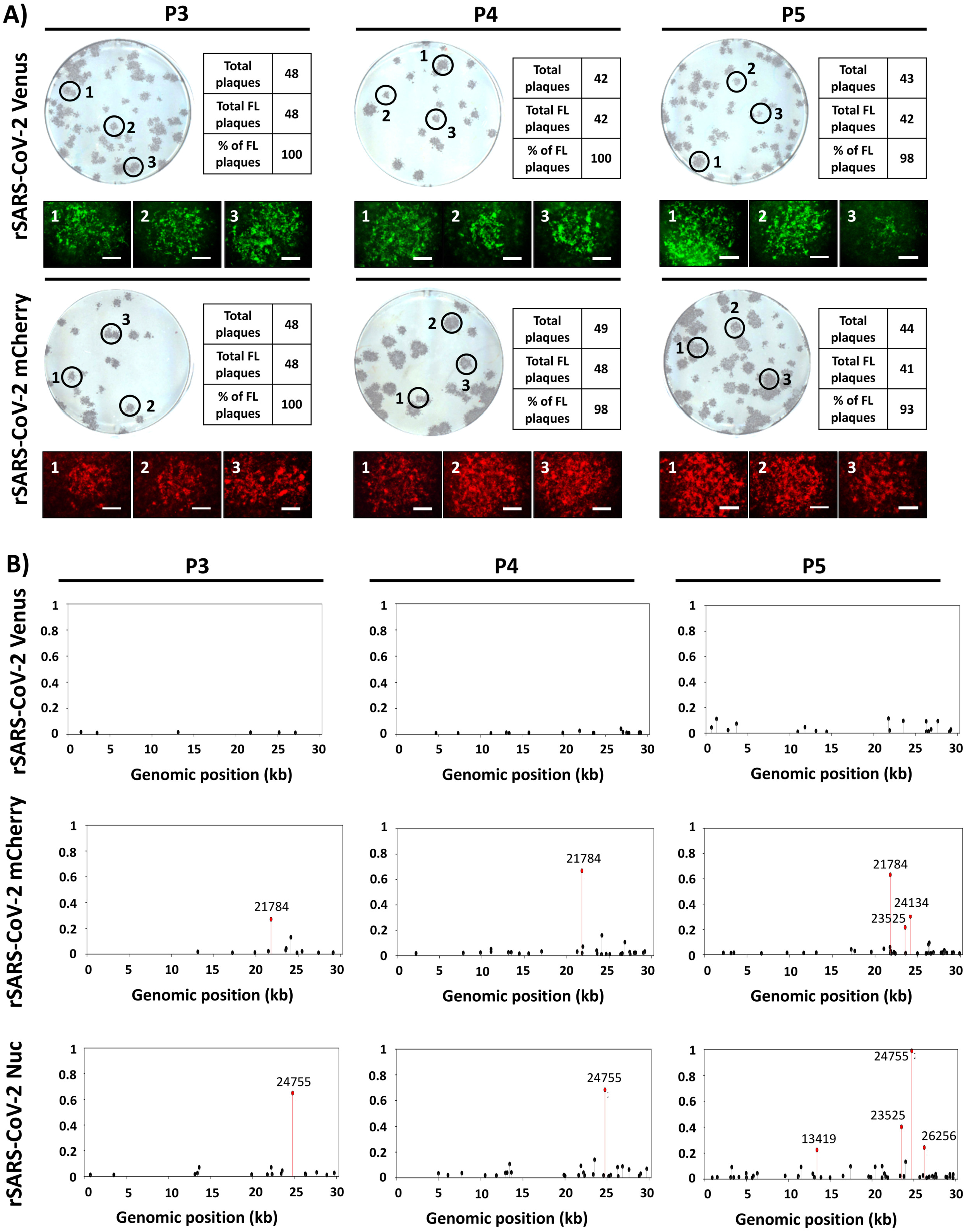
Genetic stability of fluorescent-expressing rSARS-CoV-2. **A) Plaque assay:** Fluorescent-expressing rSARS-CoV-2 were passaged up to 5 times in Vero E6 cells and infectious virus-containing tissue culture supernatants from passages 3 to 5 (P3-P5) were assessed for Venus or mCherry expression at 72 h post-infection, before immunostaining with the SARS-CoV NP MAb 1C7. The percentage of reporter-expressing viruses was determined from ∼40-50 viral plaques per passage. Representative images of immunostaining and fluorescence (4X magnification, scale bar, 750 μm) obtained from each P3-P5 viral plaques are shown. **B) Sequence analysis:** Reporter-expressing rSARS-CoV-2 non-reference allele frequencies from virus stock (P3) and after two consecutive passages in Vero cells (P4 and P5) were determined using next generation sequencing, using modified rSARS-CoV-2/WT reference genomes. Non-reference alleles that were below 1% of reads are not shown and those greater than 20% are indicated in red.

## Discussion

Reporter-expressing viruses represent a powerful tool for both basic research and translational studies (14, 15, 31–34). Several research groups, including ours, have previously described recombinant viruses expressing reporter genes to easily study the biology of viral infections, to evaluate the efficacy of antivirals or NAbs, and for *in vivo* studies in validated animal models (35–49).

Both fluorescent and luciferase proteins have been used to generate reporter-expressing viruses. However, the innate and differing properties of reporter genes dictate which one might be inserted into a recombinant virus. While fluorescent proteins provide an efficient way to track viral infections using microscopy, luciferase proteins are more readily quantifiable and therefore more amenable to HTS studies (14, 15, 50). For this reason, in this study we generated rSARS-CoV-2 expressing fluorescent (Venus and mCherry) or luciferase (Nluc) proteins (**Figure 1**). These reporter genes were selected based on either their distinctive fluorescent properties (Venus and mCherry) or because of their small size, stability, high bioluminescence activity, and ATP-independency (Nluc) (51).

Although reporter-expressing rSARS-CoV-2 similar to those reported here have been recently described (16–18), this is the first report of a replicating competent rSARS-CoV-2 expressing mCherry. Recombinant viruses expressing a red fluorescent protein represent an advantage over those expressing GFP or mNeonGreen (16–18) in that many genetically modified cell lines and/or animals express green fluorescent proteins. Another limitation of green fluorescent proteins during *in vivo* imaging is the absorption of the fluorophores’ excitation and emission by hemoglobin and autofluorescence of tissues (52–55). Recombinant viruses expressing red fluorescent proteins represent a better option to combine with genetically modified GFP-expressing cell lines and/or animals and, based on their reduced autofluorescence background, to more accurately capture the dynamics of viral infection and replication.

Reporter-expressing replicating competent viruses can be used to monitor viral infections, assess viral fitness, evaluate and/or identify antivirals and/or NAbs, where reporter gene expression can be used as a valid surrogate for viral detection in infected cells. Expression of Venus, mCherry, or Nluc from our rSARS-CoV-2 were confirmed by directly visualizing fluorescence expression under a fluorescent microscope (Venus and mCherry) or luciferase activity (Nluc) using a microplate reader (**Figures 2 and 3**). Western blot analyses using specific antibodies against each of the reporter genes further confirm expression from their respective rSARS-CoV-2 (**Figures 2 and 3**). Notably, despite deletion of the 7a ORF and insertion of a reporter gene, the three reporter-expressing rSARS-CoV-2 displayed similar growth kinetics and plaque phenotype than their WT counterpart (**Figure 3**). As expected, viral infection was visualized in real time, without the need of secondary approaches (e.g. MAbs) to detect the presence of the virus in infected cells. Overall, reporter gene expression displayed similar kinetics that correlated with levels of viral replication, further demonstrating the feasibility of using these reporter genes as a valid surrogate of assess viral infection.

Therapeutic treatment of SARS-CoV-2 infections is currently limited to the use of Remdesivir (3), and despite significant global efforts, there is no preventative vaccine for the treatment of SARS-CoV-2 infections. Notably, there is a possibility, similar to the situation with other respiratory viruses (e.g. influenza), of the emergence of drug-resistant SARS-CoV-2 variants that will impose a significant challenge to the currently ongoing COVID-19 pandemic (56). Thus, it is imperative to not only discover new antivirals and other therapeutic approaches but also prophylactics for the treatment of SARS-CoV-2 infections. To that end, rapid and sensitive screening assays to identify compounds with antiviral activity or to assess efficacy of vaccine candidates for the therapeutic and prophylactic treatment of SARS-CoV-2 infections, respectively, are urgently needed. In this study, we demonstrate that reporter-expressing rSARS-CoV-2 represent an excellent option for the rapid identification and characterization of both antivirals (**Figure 4**) and NAbs (**Figure 5**) for the therapeutic and/or prophylactic treatment of SARS-CoV-2 infections. Importantly, EC_50_ (antivirals) and NT_50_ (NAbs) obtained with our reporter-expressing viruses were comparable to those obtained using rSARS-CoV-2/WT or described by others in the literature (17, 21, 27), demonstrating the feasibility of using our reporter-based microneutralization assays for the rapid identification of antivirals or NAbs (**Figures 4 and 5,** respectively). Furthermore, our results indicate that reporter-expressing Venus, mCherry, and Nluc rSARS-CoV-2 are stable up to 5 passages *in vitro* in Vero E6 cells, including expression of the reporter gene (**Figure 6**). To date, we have not yet conducted studies to evaluate the feasibility of using these reporter-expressing rSARS-CoV-2 *in vivo*. It is possible, and similar to other respiratory viruses, that rSARS-CoV-2 expressing reporter genes could also be used to study the biology of viral infections in validated animals of viral infection.

Our SARS-CoV-2 reverse genetics based on the use of BAC have allowed us to rescue rSARS-CoV-2/WT (19) and rSARS-CoV-2 stably expressing reporter genes. In the case of our reporter-expressing rSARS-CoV-2, we removed the 7a ORF and substituted it for various reporter genes without a significant impact in viral replication. The feasibility of removing viral genes and insert reporter genes demonstrate the genetic plasticity of the SARS-CoV-2 genome and open the possibility of generating recombinant viruses expressing other genes of interest for the development of SARS-CoV-2 vaccines that could be used for the control of the currently ongoing COVID-19 pandemic.

## Acknowledgements

We want to thank Dr. Thomas Moran at the Icahn School of Medicine at Mount Sinai for providing us with the SARS-CoV cross-reactive NP MAb 1C7. We also want to thank BEI Resources for providing the SARS-CoV-2 USA-WA1/2020 isolate (NR-52281) and Marina McDew-White and Robbie Diaz for constructing the NGS libraries. Finally, we would also like to thank members at our institutes for their efforts in keeping them fully operational during the COVID-19 pandemic and the BSC and IACUC committees for reviewing our protocols in a time efficient manner. We would like to dedicate this manuscript to all COVID-19 victims and to all heroes battling this disease.

